# Vegetative Cell and Spore Proteomes of *Clostridioides difficile* show finite differences and reveal potential protein markers

**DOI:** 10.1101/598045

**Authors:** Wishwas R. Abhyankar, Linli Zheng, Stanley Brul, Chris G. de Koster, Leo J. de Koning

## Abstract

*Clostridioides difficile*-associated infection (CDI) is a health-care-associated infection mainly transmitted via highly resistant endospores from one person to the other. *In vivo*, the spores need to germinate in to cells prior to establishing an infection. Bile acids and glycine, both available in sufficient amounts inside the human host intestinal tract, serve as efficient germinants for the spores. It is therefore, for better understanding of *Clostridioides difficile* virulence, crucial to study both the cell and spore states with respect to their genetic, metabolic and proteomic composition. In the present study, mass spectrometric relative protein quantification, based on the ^14^N/^15^N peptide isotopic ratios, has led to quantification of over 700 proteins from combined spore and cell samples. The analysis has revealed that the proteome turnover between a vegetative cell and a spore for this organism is moderate. Additionally, specific cell and spore surface proteins, vegetative cell proteins CD1228, CD3301 and spore proteins CD2487, CD2434 and CD0684 are identified as potential protein markers for *C. difficile* infection.

Abstract graphic For Table of Contents Only

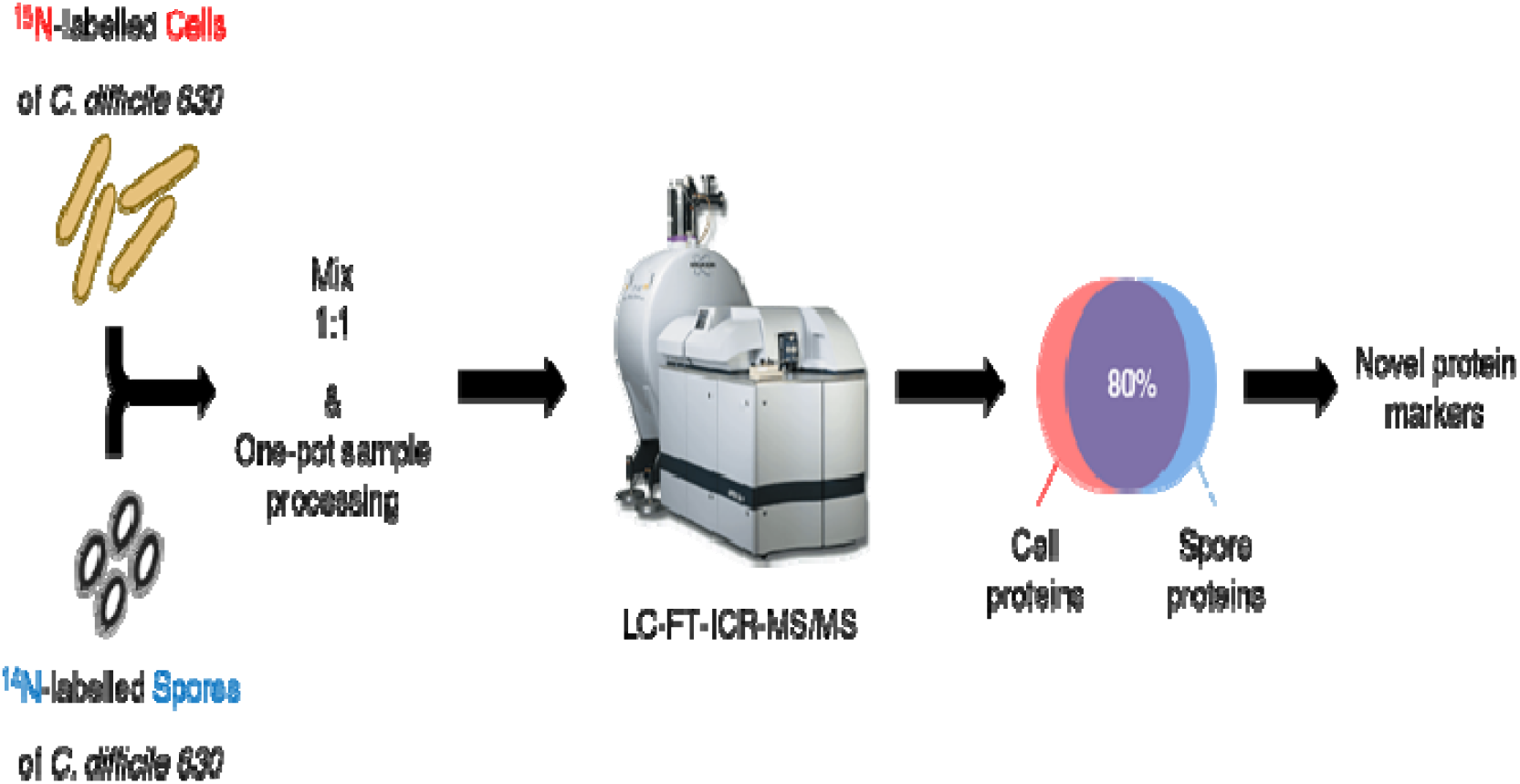

## Introduction

*Clostridioides* (previously *Clostridium*) *difficile*, an anaerobic, gram-positive pathogen, is the causative agent of an infection (CDI) characterized by pseudomembranous colitis and nosocomial diarrhoea. An over-extensive use of antibiotics is implicated for the spread of CDI and high rates of asymptomatic colonization by *C. difficile* make its diagnosis challenging (1). This necessitates designing strategies and algorithms to optimize the diagnostic tools (2). Pathogenesis of CDI is manifested via its Rho-glycosylating toxins TcdA and TcdB (3). Moreover, in response to adverse conditions, *C. difficile* can form endospores - multi-layered, highly resistant cellular entities - that are the main transmissible forms (4) in *C. difficile* infections. The spores can germinate in the intestinal environment upon interaction with bile acid mediated by the CspC protease (5), subsequently manifesting the infection.

To date, substantial research has been done on *C. difficile* vegetative cells and spores to understand the survival mechanisms of this pathogen. However, research performed on spores has gained more importance, owing to their crucial role in survival of *C. difficile*. The number of spore-related genes identified in *Clostridia* is significantly smaller than that in *Bacilli* (6). Less than 25% of the spore coat proteins of *B. subtilis* have homologues in *C. difficile* and unlike in *B. subtilis*, in *C. difficile*, neither does the activation of σ^G^ rely on σ^E^ nor is it required for the production and σ^K^ activation (7–9). Furthermore, in *C. difficile* spores, cortex hydrolysis occurs before the release of Ca^2+^-dipicolinic acid complex (10), whereas in *B. subtilis* spores, the release of this complex precedes cortex hydrolysis.

In the past decade, a few transcriptomic studies (11, 12), quantitative proteomic studies (13–19) and a lipoproteomic study (20) have been done on *C. difficile* vegetative cells. Lawley and colleagues described a protocol to purify spores and performed an extensive proteomic characterization of *C. difficile* 630 spores (21). Shortly after, we used a gel-free proteomic method and a one-pot sample processing method that focused on the spore surface layers of *B. cereus* and *C. difficile* (22). Moreover, previous studies have described exosporium removal methods for *C. difficile* spores, and examined the exosporium protein components (23) as well as spore surface glycoproteins (24). Yet, none of these studies focusses on a comparative analysis that functionally links the spore and vegetative cell proteome. In the present study, we have quantitatively characterized the *C. difficile* vegetative cell proteome relative to that of spores. To this end, spores are mixed with ^15^N-metabolically labelled vegetative cells based on spore or cell number and the mixture is processed with our recently developed one-pot method for mass spectrometric analyses, where the ^14^N/^15^N isotopic protein ratios represent the relative spore over vegetative cell protein abundances. We aim to deduce putative spore- and vegetative cell-predominant protein markers for *C. difficile*.

## Materials and Methods

### Bacterial strains, cell culture, and sporulation

*Clostridioides difficile* strain 630 (ATCC^®^ BAA1382™), acquired from the Leibniz Institute of Microorganisms and Cell Cultures, Germany, was used to derive vegetative cells and spores. All cultivations were performed at 37°C in an anaerobic chamber (Whitley DG250) supplied with a gas mixture comprising 10% hydrogen, 10% carbon dioxide, and 80% nitrogen. The cells were first grown overnight in Schaedler anaerobic broth (Oxoid, CM0497) and further passaged thrice through the newly developed ^15^N-yeastolate medium (described below) to obtain ^15^N-labelled vegetative cells. After the third passage, the cells were grown overnight until OD_600_ ≈ 1.7 and harvested by centrifugation. These cells were then aliquoted and stored at −20°C until further use. To obtain spores, the vegetative cells were pre-cultured overnight in Columbia broth and inoculated in Clospore medium (25). Typically, bottles containing 500 ml of Clospore medium, kept in the anaerobic chamber overnight, were inoculated with the pre-cultures. Spores were harvested after 2 weeks of incubation and intensively purified using a combination of ultra-sonication, enzyme treatment (lysozyme, trypsin, and proteinase K), and washing with sterile milli-Q water (21, 25). The spore crops were subjected to density gradient centrifugation by layering spores suspended in 20% Histodenz (Sigma-Aldrich, USA) on top of 50% Histodenz in 2 ml Eppendorf tubes, and centrifuging for 25 min at 15000×*g*.

### Preparation of ^15^N-yeastolate medium

*Saccharomyces cerevisiae* CEN. PK1137D was grown at 37°C in a defined CBS medium(26) modified with ^15^NH_4_Cl (replacing (NH_4_)_2_SO_4_) as the sole nitrogen source. Yeast cells were harvested by centrifugation (5000 ×*g*, 30 min) and washed with water. The protocol to generate yeastolate was adapted from previous studies (27, 28). The yeast cells were made into a 30% (w/v) slurry, ultrasonicated by a tip ultrasonicator for 30 min. The pH of the slurry was adjusted to 7.5 using NaOH before incubating under continuous shaking for 5 days at 50°C. Thereafter, the slurry was ultrasonicated again and centrifuged at 20000×*g* for 30 min to collect the supernatant. The pellet was washed twice and the supernatants were combined and lyophilized, to generate powdered yeastolate. The final yeastolate medium contained 2% ^15^N-yeastolate, 2% glucose, and 0.2% NaCl.

### One-pot sample processing

The one-pot protocol has been previously described in detail (17). Typically, spores & cells were mixed in 1:1 ratio based on the spore or cell counts and suspended in lysis buffer and disrupted for seven cycles with 0.1 mm zirconium beads (BioSpec Products, Bartlesville, OK, USA) using a Precellys 24 homogenizer (Bertin Technologies, Aix en Provence, France). The tubes were incubated for 1 h at 56°C and alkylated using 15 mM iodoacetamide for 45 min at room temperature in the dark. The reaction was quenched with 20 mM thiourea and digestion with Lys-C (at 1:200 protease/protein ratio) was carried out for 3 h at 37°C. Samples were diluted with 50 mM ammonium bicarbonate buffer and digested with trypsin (at 1:100 protease/protein ratio) was carried out at 37°C for 18 h. The tryptic digest was freeze-dried. Before use, the freeze-dried samples were re-dissolved in 0.1% TFA and desalted using Omix μC18 pipette tips (80◻μg capacity, Varian, Palo Alto, CA, USA) according to the manufacturer’s instructions.

### Fractionation of peptides

ZIC-HILIC chromatography was used to fractionate the freeze-dried peptide samples. Dried digests were dissolved in 500 μl of Buffer A (85% acetonitrile, 5 mM ammonium acetate, 0.4% acetic acid, pH 5.8), centrifuged to remove any undissolved components, and injected into the chromatography system. An isocratic flow with 100% Buffer A for 10 min was followed by a gradient of 0-30% Buffer B (30% acetonitrile, 5 mM ammonium acetate, 0.5% acetic acid, pH 3.8) in the first phase and 30-100% of Buffer B in the second phase (flow rate 400 μl/min). The peptides were eluted and collected in 10 fractions, freeze-dried, and stored at −80°C until further use.

### LC-FT-ICR MS/MS analysis

ZIC-HILIC fractions were re-dissolved in 0.1% TFA, peptide concentrations were determined by measuring absorbance at 205nm and 300ng tryptic peptide mixtures were injected for analyses. LC-MS/MS data of each ZIC-HILIC fraction were acquired with an Apex Ultra Fourier transform ion cyclotron resonance mass spectrometer (Bruker Daltonics, Bremen, Germany) equipped with a 7 T magnet and a Nano electrospray Apollo II Dual Source coupled to an Ultimate 3000 (Dionex, Sunnyvale, CA, USA) HPLC system. LC conditions and acquisition parameters were as described previously (17).

### Data analysis and bioinformatics

Each raw FT-MS/MS data set was mass calibrated better than 1.5 ppm on the peptide fragments from the co-injected GluFib calibrant. The 10 ZIC-HILIC fractions were jointly processed as a multifile with the MASCOT DISTILLER program (version 2.4.3.1, 64 bits), MDRO 2.4.3.0 (MATRIX science, London, UK), including the Search toolbox and the Quantification toolbox. Peak-picking for both MS and MS/MS spectra was optimized for the mass resolution of up to 60000. Peaks were fitted to a simulated isotope distribution with a correlation threshold of 0.7, with minimum signal to noise ratio of 2. The processed data were searched in a MudPIT approach with the MASCOT server program 2.3.02 (MATRIX science, London, UK) against the *C. difficile* 630 ORF translation database. The MASCOT search parameters were as follows: enzyme - trypsin, allowance of two missed cleavages, fixed modification - carboamidomethylation of cysteine, variable modifications - oxidation of methionine and deamidation of asparagine and glutamine, quantification method - metabolic ^15^N labelling, peptide mass tolerance and peptide fragment mass tolerance - 50 ppm. MASCOT MudPIT peptide identification threshold score of 20 and FDR of 2% were set to export the reports.

Using the quantification toolbox, the quantification of the light spore peptides relative to the corresponding heavy cell peptides was determined as light/heavy ratio using Simpson’s integration of the peptide MS chromatographic profiles for all detected charge states. The quantification parameters were: Correlation threshold for isotopic distribution fit - 0.98, ^15^N label content - 99.6%, XIC threshold - 0.1, all charge states on, max XIC width −120 seconds, elution time shift for heavy and light peptides - 20 seconds. All isotope ratios were manually validated by inspecting the MS spectral data. The protein isotopic ratios were then calculated as the average over the corresponding peptide ratios. For each of the three replicas, the identification and quantification reports were imported into a custom made program to facilitate data combination and statistical analysis. Protein identification was validated with identifications in at least two replicas. For these identified proteins the relative quantification was calculated as the geometric mean of at least two validated light/ heavy ratios. All identification and quantification protein data are listed in **Table S1**. The mass spectrometry proteomics data have been deposited as a partial submission to the ProteomeXchange Consortium via the PRIDE (29) partner repository with the dataset identifier PXD012030.

Transmembrane proteins were predicted using the default parameters on the TMHMM Server (version 2.0 http://www.cbs.dtu.dk/services/TMHMM/). DAVID Bioinformatics Resources tool (version 6.8) was used (30) to retrieve the functional annotation data of UniProt key word and KEGG pathway classifications. The BioCyc pathway analysis tool (31) was used to generate a cellular overview of the quantified proteins.

## Results

### Metabolic labelling of *C. difficile* vegetative cells using ^15^N-yeastolate

As illustrated in **Fig. 1** our culturing methods successfully yielded ^15^N labelled vegetative cells and ^14^N spores. For a number of identified ^15^N labelled vegetative cell peptides, the ^15^N label content has been calculated based on their mass spectrometric isotope patterns using the NIST isotope calculator (32). This shows that the present metabolic labelling method achieves a ^15^N label content of ≥ of 99.5%, which is amply sufficient for accurate protein quantification.

**Figure 1.**
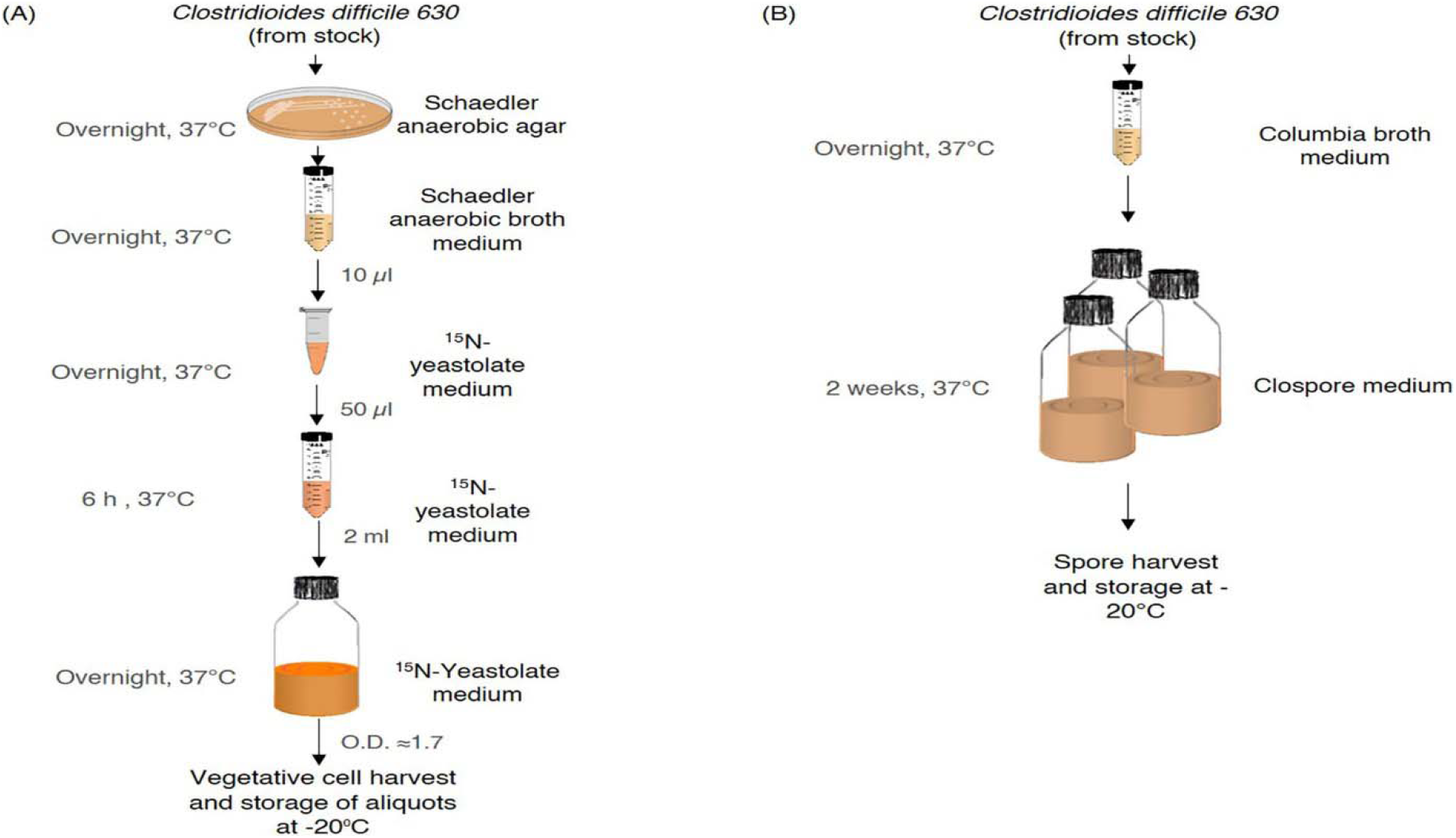
Preparation workflow of (A) ^15^N-labelled vegetative cells and (B) ^14^N spores of *Clostridioides difficile* 630. See the Materials and Methods section for more details. The images for petri dish (http://www.clker.com/clipart-red-petri-dish-3.html), media bottle (http://www.clker.com/clipart-reagent-bottle-with-growth-media.html), the Eppendorf tube (https://www.clipartmax.com/middle/m2i8H7m2A0G6N4G6_isop-eppi-pellet-zymo-clip-art-at-clker-eppendorf-tube/) and 50 ml tube (https://openclipart.org/detail/170165/50ml-centrifuge-tube) are obtained from copyright-free public domain websites and further modified using Microsoft Power Point 2016.

### Identification and quantification of cell and spore proteins

A total of 1095 proteins has been identified from *C. difficile* spores and vegetative cells of which 796 have been relatively and reproducibly quantified between spores and vegetative cells (**Supplementary Table S1**). **Fig. 2** represents a distribution of quantified proteins, where the abundance of the combined spore and vegetative cell proteins indicated by the log_10_ values of their MASCOT scores are plotted against the corresponding relative protein levels in the two morphotypes indicated by the log_2_ values of the light/heavy ratios. Eighty seven proteins are considered to be predominant present in spores with a light/heavy ratio > 20, while 81 proteins are considered to be predominant present in vegetative cells with light/heavy ratios < 0.05. From the remaining 628 commonly shared proteins 18% are enriched in spores with light/heavy ratio between 1 and 20 while 82% are enriched in cells with a light/ heavy ratio between 1 and 0.05. In total, 167 proteins have been classified as membrane proteins by the TMHMM analysis (**Supplementary Table S2**). The cellular overview based on pathway analysis of the quantified proteins is represented in **Supplementary Fig. S1** indicating the pathways to which these commonly shared proteins belong.

**Figure 2.**
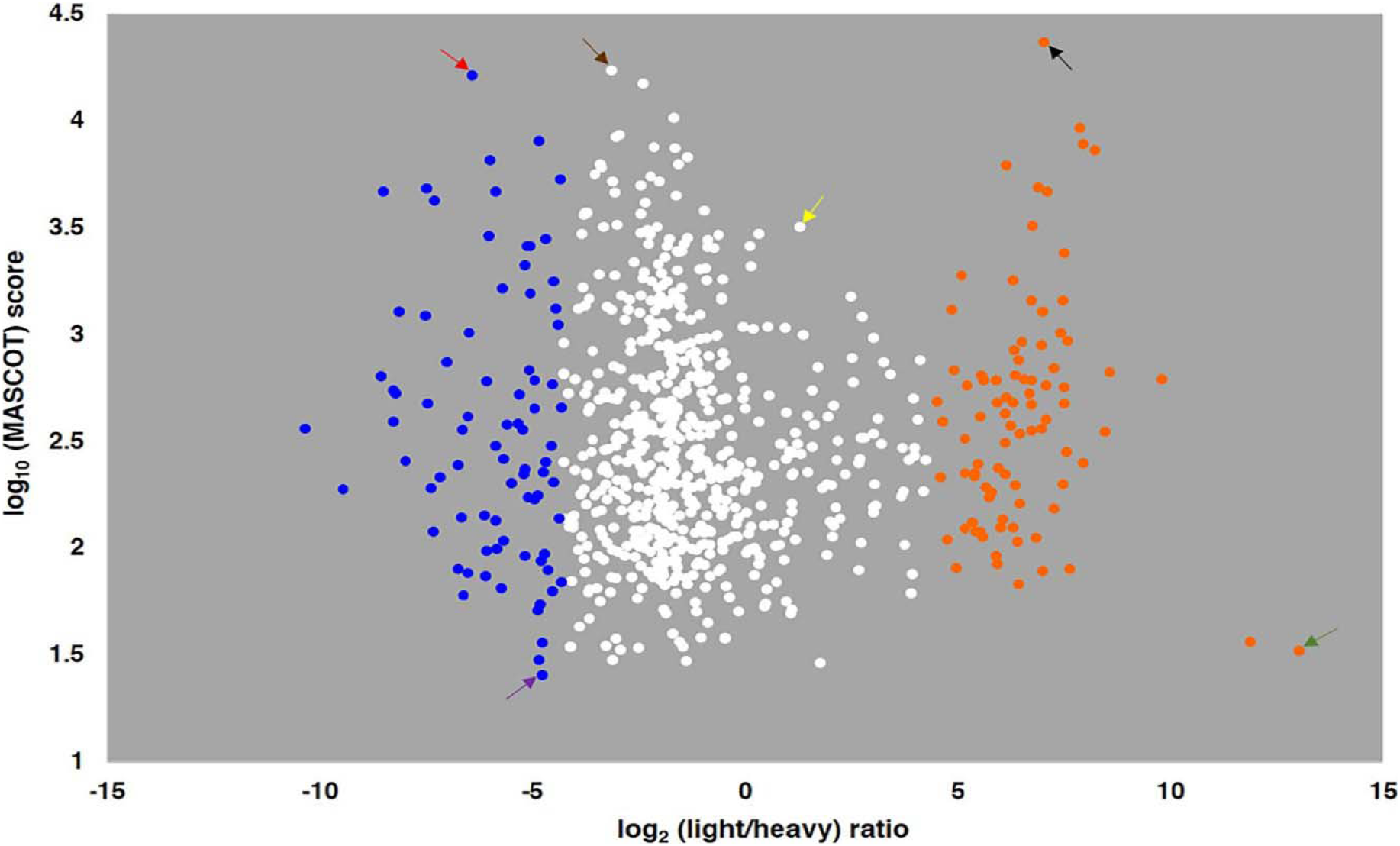
Distribution of proteins in *Clostridioides difficile* 630 spores and vegetative cells. MASCOT score indicates the combined spore and cell abundance of a protein versus its light/heavy protein isotopic ratio, which represents the relative level of the protein in spores over vegetative cells. The orange dots indicate spore-predominant proteins (light/heavy ratios > 20), blue dots indicate vegetative cell-predominant proteins (light/heavy ratios < 0.05), and white dots indicate proteins common between spores and vegetative cells (20> light/heavy ratios >0.05). Black arrow - SspA; green arrow- CD2657; red arrow - SlpA; purple arrow- CD0594; brown arrow - CD0825; and yellow arrow - CD0718. (See the text and **Table S1** for more details).

### Spore-predominant proteins

These include proteins from the spore coat and exosporium layers, classified under UniProt Keywords virion and capsid proteins, along with some rotamase proteins and metalloproteases (**Table 1**). SspA is the most abundant protein in this category, whereas an uncharacterized protein CD2657 (with 13 times higher levels in spores than in vegetative cells) is the least abundant (**Fig. 2**). The TMHMM analysis classified 26 proteins from this category as membrane proteins (**Supplementary Table S2**). Most of these are uncharacterized membrane proteins but some are known proteins such as SpoVD, SpoVAC, SpoVFB, FtsH, and DacF.

**Table 1.** Uniprot keywords annotation enrichment of quantified *Clostridioides difficile* 630 spore- and vegetative cell proteins based on DAVID functional annotation analysis.

| UniProt Keyword<sup>a</sup> | No. of proteins |  |  |
| --- | --- | --- | --- |
|  | Spore<br>proteome | Vegetative cell<br>proteome | Commonly<br>shared |
| Aminotransferase |  | 10 |  |
| Arginine biosynthesis |  | 7 |  |
| Cell shape |  | 8 |  |
| Elongation factor |  | 5 |  |
| Peptidoglycan synthesis |  | 7 |  |
| Cytoplasm | 130 | 136 | 127 |
| Transferase | 115 | 121 | 110 |
| Nucleotide-binding | 107 | 114 | 105 |
| Hydrolase | 107 | 104 | 94 |
| ATP-binding | 89 | 95 | 87 |
| Metal-binding | 82 | 80 | 78 |
| Oxidoreductase | 61 | 65 | 54 |
| Ribonucleoprotein | 48 | 48 | 48 |
| RNA-binding | 49 | 48 | 48 |
| Ligase | 46 | 48 | 46 |
| Ribosomal protein | 47 | 47 | 47 |
| Protein biosynthesis | 34 | 36 | 34 |
| Lyase | 31 | 36 | 28 |
| Magnesium | 33 | 33 | 33 |
| rRNA-binding | 32 | 32 | 32 |
| Zinc | 30 | 28 | 28 |
| Amino-acid biosynthesis | 22 | 27 | 22 |
| Isomerase | 25 | 26 | 24 |
| Aminoacyl-tRNA synthetase | 23 | 24 | 23 |
| Protease | 25 | 20 | 20 |
| GTP-binding | 15 | 16 | 15 |
| Pyridoxal phosphate | 13 | 14 | 12 |
| Glycosyltransferase | 14 | 13 | 13 |
| Cell cycle | 11 | 13 | 11 |
| Cell division | 11 | 13 | 11 |
| Flavoprotein | 14 | 12 | 11 |
| FAD | 11 | 10 | 9 |
| NADP | 11 | 12 | 11 |
| tRNA-binding | 11 | 11 | 11 |
| NAD | 12 | 11 | 10 |
| Ion transport | 10 | 11 | 10 |
| Pyruvate | 11 | 11 | 11 |
| Chaperone | 10 | 10 | 10 |
| Manganese | 10 | 9 | 9 |
| ATP synthesis | 9 | 9 | 9 |
| Aminopeptidase | 9 | 9 | 9 |
| Lysine biosynthesis | 8 | 8 | 8 |
| Hydrogen ion transport | 8 | 8 | 8 |
| Glycolysis | 8 | 7 | 7 |
| Stress response | 7 | 7 | 7 |
| Diaminopimelate biosynthesis | 6 | 6 | 6 |
| Pyrimidine biosynthesis | 6 | 6 | 6 |
| CF(1) | 5 | 5 | 5 |
| One-carbon metabolism | 5 | 5 | 5 |
| DNA-directed RNA polymerase | 4 | 4 | 4 |
| Methionine biosynthesis | 4 | 4 | 4 |
| Virion | 10 |  |  |
| Capsid protein | 9 |  |  |
| Rotamase | 5 |  |  |
| Metalloprotease | 6 |  |  |

<sup>a</sup> EASE score i.e. *p*-value threshold for the keyword annotation enrichment was set to 0.05

### Cell-predominant proteins

These include the cytoplasmic proteins such as aminotransferases, arginine biosynthesis, elongation factors, cell shape and peptidoglycan synthesis proteins (**Table 1**). The surface layer protein SlpA is the most abundant and unique protein in this category, whereas an uncharacterized protein CD0594 (with levels 27 times higher in vegetative cells than in spores) is the least abundant (**Fig. 2**). Eighteen membrane proteins are predominant in vegetative cells, as predicted by TMHMM (**Supplementary Table S2**).

### Proteins shared between spores and vegetative cells

The proteins shared between spores and cells are mostly ribosomal proteins, cell cycle-regulating and/or associated proteins, and cytosolic proteins involved in pathways required for anabolism and catabolic pathways of energy metabolism distributed over 46 categories by DAVID (**Table 1**, **Supplementary Fig. S1**). These also include products of 25 essential genes(33) such as peptidoglycan synthesis protein MurG (CD2651) and formate-tetrahydrofolate ligase CD0718 (2 and 2.4 times higher levels in spores than in vegetative cells, respectively), and S-adenosylmethionine synthase MetK as well as a rubrerythrin CD0825 (~4 and ~9 times higher levels in vegetative cells than in spores, respectively) (**Fig 2**). The TMHMM analysis of the shared proteins identified 123 membrane proteins (**Supplementary Table S2**), such as the phosphotransferase system (PTS) of sugar transporters, ABC-type transporters, and V-type ATPases. These also included most proteins involved in the Wood-Ljungdahl pathway (**Fig. 3**). From **Table 1** it can be deduced that proteins from this category that are present in spores are essentially those that are required for hibernation, the initiation of growth and the resumption of metabolism upon germination and initiation of outgrowth.

**Figure 3.**
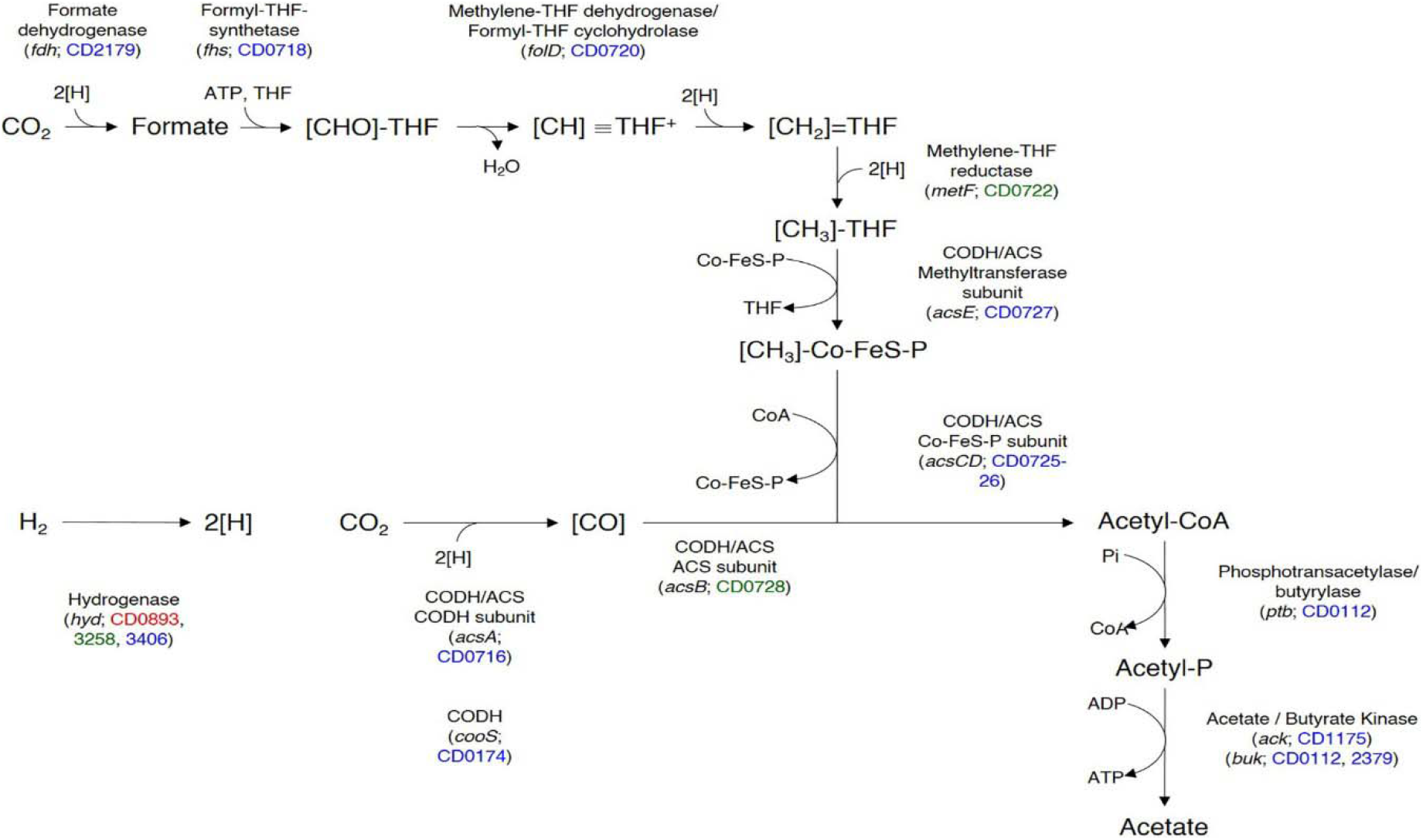
Classification of proteins associated with the Wood-Ljungdahl pathway identified in *Clostridioides difficile* 630. Proteins presented in red, green, and blue fall under the categories of vegetative cell-predominant, spore-predominant, and shared proteins, respectively.

## Discussion

To our knowledge, the proteomes of *C. difficile* vegetative cells and spores have been explored for the first time in a single experimental set-up to understand the fundamental differences between these two morphological forms of the bacterium. To this end, spores and ^15^N-labelled vegetative cells have been mixed for relative quantification. Metabolic labelling using ^15^N isotopes is a highly accurate means of proteome quantification (34). However, a method for labelling *C. difficile* was unavailable until recently (19). Here a method for metabolic labelling using ^15^N-labelled yeastolate medium has been developed which provides a simple, economical, and rapid means to perform quantitative proteomics of a variety of pathogenic and non-pathogenic microbes. Our analyses show that 80% of the quantified proteins are common for both cells and spores, indicating that pathogenic *C. difficile* employs a relatively modest proteomic changeover to enable long-term survival as a dormant spore. The corresponding pathways are shared between vegetative cells and spores (**Supplementary Fig. S1**) however the relative quantities of these proteins vary between cells and spores. A discussion of quantified and functionally key proteins of *C. difficile* is presented below.

*Clostridioides difficile* expresses an array of cell surface proteins, including the S-layer protein SlpA and its paralogues from the cell wall protein (CWP) family. Eight Cwp family proteins are quantified: Cwp18 and 22 are identified in both morphotypes, with higher levels in spores, whereas Cwp2, 5, 6, 19, 84, and CwpV are identified in vegetative cells. Cwp22, a functional homologue of LD-transpeptidase (Ldt_cd2_), is an important spore protein that plays a role in peptidoglycan remodelling and confers resistance to β-lactam antibiotics (35, 36). CwpV promotes *C. difficile* aggregation and its strain-dependent structural variations may assist in evading the host antibody response (37) or to launch an anti-phage strategy (38). *Clostridioides difficile* spores frequently have an interspace region between the spore coat and the fragile, heterogeneous exosporium (39, 40). Although most known and putative exosporium proteins described previously (23) have been identified in this study, the BclA family of proteins have not (except BclA1 (CD0332), identified in one replicate and thus not quantified). An absence of hair-like structures in the *C. difficile* 630 exosporium (40) may underlie this finding. Other identified proteins such as CD1474, CD2845, and CD1524 - all rubrerythrins -likely present in the exosporium, may play a role in fighting reactive oxygen species and oxidative stress (41).

*Clostridioides difficile* relies heavily on the phosphoenolpyruvate-dependent phospho-transferase system (PTS) for uptake and regulation of various sugars and sugar derivatives (42). The PTS is clearly advantageous for germinating spores, since various sugars are readily available in the human gut and may thus be used to facilitate outgrowth and establish infection (43, 44). We have identified 22 PTS proteins, of which 15 could be quantified (**Supplementary Table S1**). The quantified PTS proteins, except CD3027, are shared between vegetative cells and spores. Along with the central PTS proteins HPr (PtsH/CD2756) and Enzyme I (PtsI/CD2755), these proteins function in the transport of fructose (FruABC/CD2269, CD2486-88), glucose (PtsG/CD2667, CD3027, CD3089), mannitol (MtlF/CD2332 and MtlA/CD2334)), mannose (CD3013-14), xyloside (XynB/CD3068), and β-glucoside (BglF5/CD3137). Although shared, CD3013-14, CD2486-88, CD3089, PtsH, and MtlA-F showed relatively higher levels in spores, whereas PtsG, PtsI, BglF5, FruABC, CD3027, and XynB show lower levels in spores. Interestingly, a previous study has shown that in germinating spores, *bglF5* and *ptsG* transcripts are downregulated whereas those of *fruABC* and *cd2486*-*87* are upregulated (45). In mouse infections, *C. difficile* CD2487 is upregulated 14 h post-infection whereas proteins XynB, CD3027, and PtsI are downregulated 38 h post-infection (11). In pig infections of *C. difficile* PtsI, BglF5 (4-12 h post-infection) and MtlA (only 12 h post-infection) are upregulated (46) and XynB and CD3013-14 are downregulated. Furthermore, MtlA and MtlF can repress *tcdA* and *tcdB* toxin expression in *C. difficile* (47). Put together, these studies indicate that PtsI, BglF5, CD2487, and MtlA potentially play a role in the pathogenesis of *C. difficile* infections; CD2487 and MtlA are predominantly present in spores, making them important targets in understanding spore persistence.

Non-PTS transport systems are also involved in carbohydrate uptake in clostridia (42). Of all ATPases and related proteins quantified, only 6 and 4 proteins belong to the cell-predominant and spore-predominant categories, respectively. Four V-type ATPases have been quantified; however, only AtpC is spore-predominant whereas the other three are also expressed in cells. AtpC is associated with proton transport, possesses a hydrolase activity, and contains a CodY-binding region (48) potentially repressing toxin expression in *C. difficile* (49). It also regulates synthesis and circulation of pyruvate and 2-oxoglutarate in the cell (50), providing proteomic flexibility during spore revival. The spore-predominant ATPases include a cation (Ca^2+^)-transporting ATPase (CD2503), which shares 42% identity with *B. subtilis* YloB ATPase, likely responsible for accumulating intracellular calcium and reinforcing thermal resistance(51). Two ABC-type transporters - lipoproteins CD2365 and SsuA - are expressed in both *C. difficile* vegetative cells and spores (20) but are present at higher levels in spores. These are alkanesulfonate and taurine binding proteins, respectively and SsuA is involved in sulfur metabolism (52). Interestingly, the taurine side chain of taurocholate selectively binds its potential receptor site(53) and taurine itself is an alkane sulfonate, thus higher levels of SsuA and CD2365 in spores could indicate potential taurine interaction during germination, similar to CD3298(54). Identified protein CD0114 shares 25% identity with CD3298, making it worth studying for a possible role in spore germination. Additionally, CD3669 - an orthologue of GerM -might be involved in sporulation (55).

The ‘stay-green’ family protein CD3613 is a putative exosporium protein. Usually, these proteins perform chlorophyll degradation (56) but the upregulation of CD3613 during sporulation in a mouse model (11) suggests a potential role in pathogenesis. CD2434 has an UBA_NAC_like bacterial protein domain commonly found in proteins involved in ubiquitin-dependent proteolysis (57). Although not a direct evidence, this observation suggests involvement of CD2434 in pathogenesis; a previous study has shown that the *E. coli* toxin CNF1 utilizes the ubiquitin-proteasome assembly of host cells to partially inactivate their Rho GTPases (58), a mechanism similar to that of TcdA and TcdB toxins (59). The spore envelope protein CD2635 may be involved in germination (12). This protein, similar to CD2636, contains a characteristic YIEGIA domain and both could play significant roles in spore assembly as well as disintegration.

Amino acids play a crucial role in spore germination and functioning of the Stickland pathways in *C. difficile*. CD3458 and CD1555 are putative amino acid permeases identified to be slightly more abundant in spores than in vegetative cells. CD3458 contains a putative amino acid permease domain and an SLC5-6-like_sbd superfamily domain, thus qualifying as an amino acid permease and sodium/glucose co-transporter. Another amino acid permease CD2612, identified in vegetative cells, is upregulated in the presence of cysteine (52), implying a role in sulfur metabolism. CD2344 contains an Asp-Al_Ex domain found in aspartate-alanine antiporters and might be capable of developing a membrane potential enough to carry ATP synthesis via FoF1 ATPase (60).

Peptidases and proteases are crucial for vegetative cell and spore survival. In this study, 29 peptidases and 12 proteases are identified and quantified. These include proteins involved in germination, such as Gpr (CD2470) and CspBA. In the present study, CspC protease has been detected in only one replicate and thus not quantified. Other proteins involved in cellular regulatory processes, such as ATP-dependent Clp proteases, zinc metalloproteases, serine proteases, Lon proteases, have also been quantified in the present study. These peptidases belong to various families such as aminopeptidase (M1), metalloendopeptidase (M16), aminopeptidase (M18), membrane dipeptidase (M19), glutamate carboxypeptidase (M20), glycoprotease (M22), methionyl aminopeptidase (M24), prolyl oligopeptidase (S9), and collagenase (U32). It is speculated that proteins belonging to the M22 and U32 family (CD0150 and CD1228, respectively) function in spore germination. In fact, BA0261, a CD0150 orthologue in *B. anthracis*, is suggested to play a role in spore germination (61) and a collagenase is implicated in virulence of *B. cereus* endophthalmitis (62) indicating a similar potential for CD1228. A previous study suggested that a Lon protease in *Brucella* sp. is involved in BALB/c mice infections (63). Thus, the Lon protease CD3301, present in vegetative cells and spores, also likely plays a role in infection. CD0684 has been previously reported to be present in *C. difficile* spores under σ^G^ regulation and suggested to be involved in stress resistance (12). Notably, in the present study, none of the identified peptidases or proteases are found to be predominant in vegetative cells.

From the quantified dataset 198 proteins are classified to the metabolic pathways category by DAVID analysis. Although the spores are metabolically dormant, the proteins belonging to the amino acid biosynthesis, purine metabolism, glycolysis, fatty acid metabolism, and nitrogen metabolism are present and form the core protein set in spores. Moreover, in spores, arginine biosynthesis pathway proteins are present at ~20% of the levels detected in vegetative cells. This indicates that germinating spores require *de novo* synthesis of these proteins post-germination to assist the outgrowing cells. Ribosomal proteins - except the 50S ribosomal protein L30 (CD0881) - are also present in low amounts in *C. difficile* spores. CD0881 has a ferredoxin-like fold, resembling the structure of yeast L7 proteins, and is likely involved in processing precursors of large rRNAs (64), a function that could well aid the outgrowing spores. The phosphate butyryltransferases (CD0715 and CD0112/Ptb) involved in the butanoate metabolism pathway are present not only in vegetative cells but also in spores, thus conferring on the spores a metabolic flexibility.

*Clostridioides difficile* may deploy several sulfur and nitrogen metabolism proteins while surviving in anaerobic conditions. Of these, only CD2431, a nitrite/sulfite reductase, is abundant in spores. This protein also contains a 4Fe-4S domain and can catalyse the reduction of sulfite to sulfide and nitrite to ammonia (65). *Clostridioides difficile* and other acetogenic clostridia have acquired such metabolic flexibility that they can directly utilize the CO_2_ and H_2_ from air and yield a variety of products including acetate and methane (66). The Wood-Ljungdahl pathway of acetogenesis is believed to be the first biochemical pathway to have emerged on earth (67) and all proteins involved in this pathway are identified in *C. difficile* 630, which reinforces the acetogenic nature of *C. difficile* growth. Of these, CD3405, CD3407 and CD0730 have been detected only in single replicates and thus are not quantified. The other Wood-Ljungdahl pathway proteins have all been quantified, with only three proteins - MetF (CD0722), CD0728, and CD3258 - being highly abundant in spores. In contrast, only a single protein - CD0893 - is predominant in vegetative cells.

The acetogenic mode of life of *C. difficile* requires specific enzymes, such as acetyl-CoA synthases/CO dehydrogenases (CD0174, CD0176, and CD0727), formate dehydrogenases (CD2179), and iron-only hydrogenases (CD0893, CD3258, and CD3406). Enzymes CD0174 and CD0176 synthesize the key metabolite acetyl-CoA from CO, methyl corrinoid, and CoASH. The formate dehydrogenases can be seleno (CD3317) or non-seleno (CD0769 and CD2179) enzymes. Protein CD2179, an anaerobic dehydrogenase, reduces CO_2_ to formate which is further metabolized to acetyl-CoA through enzymatic reactions, one of them involving another acetyl-CoA synthase/CO dehydrogenase with a methyltransferase subunit. In acetogens lacking cytochromes, the Rnf complex (encoded by CD1137-42) is the putative coupling site for energy conservation (66). In the present study, all components of the Rnf complex, except CD1140-41, have been identified. The Rnf complex proteins, together with electron transport flavoproteins etfA2/B2 (CD1055-56), are employed in butyrate formation (68). However, the present study has identified only etfA1/B1 and etfA3/B3 proteins. These proteins are predominant in vegetative cells, indicating that they likely function exclusively during the vegetative life cycle of *C. difficile*.

The iron-only hydrogenases, 10 times more efficient in hydrogen production than [NiFe] hydrogenases (69), are abundant in clostridia. *Clostridioides difficile* encodes two trimeric and three monomeric hydrogenases (70). Moreover, proteins CD3405-3407 function as electron-bifurcating hydrogenases whereby physiological electron carriers such as ferredoxin are used for H_2_ production (71). In the present study, CD3258 is seen predominantly in spores whereas CD0893 occurs mostly in vegetative cells. Both proteins are monomeric, ferredoxin dependent(71), and contain a H-cluster i.e. a centre for hydrogen production (72). However, CD3258 has a sequence of eight cysteines for stabilizing two [4Fe4S] clusters transferring electrons from the surface to the protein’s active site (73) whereas CD0893 has a single FeS domain with a (Cx_1-4_Cx_5-9_Cx_3_C) arrangement at its N-terminus and the H-cluster has an additional cysteine residue (TSCCCPxW(70)). The predominant expression of CD3258 and CD0893 in spores and vegetative cells, respectively, indicates the distinct roles of these proteins in *C. difficile* physiology.

In addition, a few noteworthy and yet uncharacterized proteins are detected in our study. CD1470, a sulfotransferase, may be involved in cyanide detoxification. PdaA1 (CD1430) and PdaA2 (CD2719) have recently been shown to be important for cortex muramic acid-δ-lactam synthesis; spores lacking it are heat sensitive, deficient in germination, and exhibit late virulence (74). CD2719 is not identified in the present study; however, CD1556, an orthologue of PdaA2 of *B. cereus* var. *anthracis* is identified. Thus, CD1556 may be important for spore structure and germination. CD1319, an orthologue of YlxY, of *B. subtilis* (75) may be important for sporulation.

## Conclusions

The one-pot sample processing method along with ^15^N metabolic labelling has enabled a reproducible, combined cell and spore quantitative proteome analysis of the anaerobic pathogen *C. difficile* 630. The analysis outlines a relatively modest proteomic adaptation of this evolutionarily and clinically important anaerobic pathogen, when as a survival strategy, it completes spore formation. In addition to specific cell and spore surface proteins, the study has qualified vegetative cell proteins CD1228, CD3301 and spore proteins CD2487, CD2434 and CD0684 as potential protein markers for *C. difficile* infections.

## Acknowledgements

W.R.A is supported by the grant NWO ALWOP.260. L.Z. acknowledges the Erasmus Mundus program (EMEA3) and TNO (Healthy Living) for funding this research.

## Author Contributions

W.R.A analysed the data, prepared the figures and tables and wrote the main manuscript text. L.Z. performed the experiments. L. dK conceptualized and designed the experiments as well as curated and processed the proteomics data. S.B. and C. dK supervised and mentored the research. All authors reviewed the manuscript.

## Competing interests

The authors declare no competing interests.

## Supporting Information

**Supplementary Figure S1.**
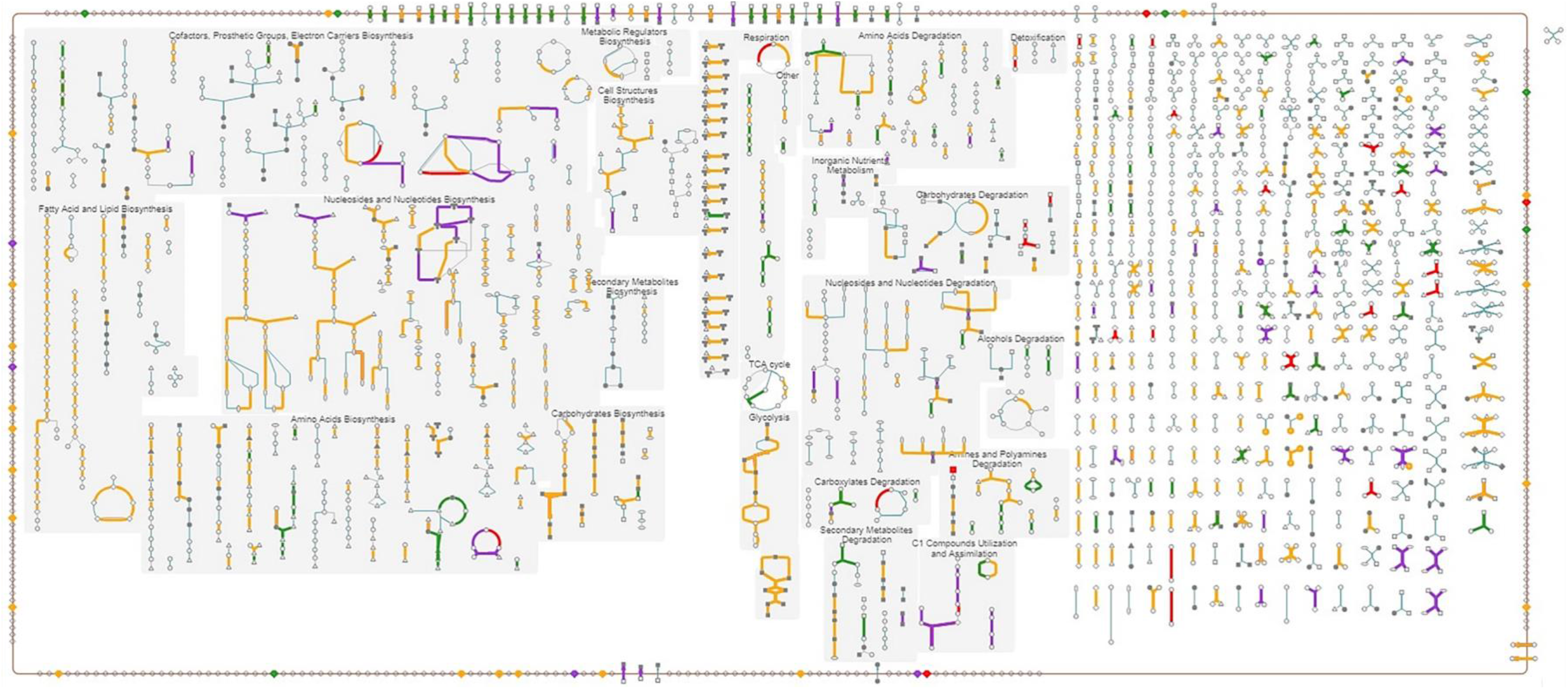
Cellular overview of quantified proteins from spores and vegetative cells of *Clostridioides difficile* 630. The overview was generated using the BioCyc pathway analysis tool. The quantified proteins that are spore-predominant (red), commonly shared but still higher in spores (purple), commonly shared but higher in cells (orange) and cell-predominant proteins (green) are represented with the pathways to which they belong. Refer to **Supplementary Table S1** for the details.

**Supplementary Table S1. Proteins identified and quantified from *Clostridioides difficile* 630 vegetative cells and spores.** Score (AA): Arithmetic average of Mascot scores from three replicates; SEM (AA): Standard Error of Means of arithmetic averages; L/H (GM): Geometric means of Light/Heavy ratios from three replicates; SEM (AA): Standard error of means in arithmetic averages of Light/Heavy ratios; SD (GM): Geometric standard deviation in Light/Heavy ratios; Proteins identified only in a single replicate are shown in red; Proteins encoded by essential genes are shown in blue.

**Supplementary Table S2. Predicted membrane proteins from *Clostridioides difficile* 630 vegetative cells and spores.** All the default parameters were used for TMHMM predictions.

## Notes

#### Summary of Updates

This version of the manuscript has been revised to update the scientific nomenclature of the organism involved. The name *Peptoclostridium difficile* has been corrected to *Clostridioides difficile* throughout the article including the supplementary files.

